# Mitigating the Effects of Population Stratification in Gene-Gene Interaction Studies

**DOI:** 10.64898/2026.08.18.745398

**Authors:** Neelotpal Das, Masao Ueki, the Alzheimer’s Disease Neuroimaging Initiative

## Abstract

Population stratification is a major source of inflated false positive rates in genome wide association studies. However, relatively few studies have examined its impact on gene-gene interaction detection, despite the importance of epistasis for understanding the genetic architecture of complex traits. In this study, we identify scenarios under which population stratification can inflate the interaction test statistics. Through analytical derivations and simulation studies, we show that this inflation is not adequately controlled by including principal components as covariates in the regression model. We then propose an alternative approach that effectively controls the inflation of false-positive rates for interaction test statistics due to population stratification by using single nucleotide polymorphism-by-population structure interaction as an additional covariate term in the regression model.

## 1 Introduction

One of the central goals of human genetic studies is to characterize the genetic architecture of complex diseases. Toward this goal, genome-wide association studies (GWAS) have identified thousands of common genetic variants, including single nucleotide polymorphisms (SNPs) and insertions/deletions (indels), associated with a broad range of complex traits and diseases (Abdellaoui et al., 2023). Causal variants however rarely act in isolation to manifest a disease, necessitating a deeper understanding of gene-gene interactions, also known as epistasis (Cordell, 2009; Goudey et al., 2017). The statistical definition of epistasis, as originally described by R.A. Fisher, is the non-random departure from a purely additivity between two or more loci (Cordell, 2002). Interactions are usually detected by testing the interaction term within a regression framework.

While performing GWAS, researchers frequently encounter the challenge of population stratification, a phenomenon characterized by the presence of latent subgroups with distinct ancestral backgrounds. This structure typically leads to the inflation of association test statistics, resulting in an increased false positive rate and a loss of power to detect true associations (Hellwege et al., 2017). To mitigate these effects, several robust methodologies are commonly employed, including genomic control (Devlin & Roeder, 1999), principal component-based EIGENSTRAT (Price et al., 2006), and linear mixed-model frameworks such as EMMAX (Kang et al., 2010) and fastGWA (Jiang et al., 2019). Although extensive focus has been dedicated to addressing population stratification within standard GWAS, this issue has received comparatively limited attention in the context of epistasis detection. There has been no study to measure the effects of population stratification on interaction test statistics under the simple regression framework. Several methods based on linear mixed models (Ning et al., 2018; Tyler et al., 2021) and multifactor dimensionality reduction (MDR) (Abegaz et al., 2021; Niu et al., 2011) have been developed to address this problem. Nevertheless, such approaches can be computationally demanding and resource intensive. In addition, MDR does not explicitly parameterize SNP main effects and may be less effective than regression-based methods in controlling false positive rates.

In this study, we investigate the impact of population stratification on epistasis detection under three regression models. The first model follows the standard regression-based definition of interaction by including a multiplicative SNP-SNP interaction term. The second model extends the first by adjusting for population structure using principal components, as in the EIGENSTRAT framework. Because the second model did not fully control inflation in interaction test statistics, we developed a third model, which is an application of the approach proposed by An et al. (2019), which additionally includes SNP-by-principal-component interaction terms.

Through analytical derivations, we demonstrate that the proposed third model reduces false-positive inflation in SNP–SNP interaction testing. We further evaluate its performance through simulation studies involving both continuous and binary phenotypes, showing that the proposed approach provides improved control of false-positive rates under population stratification. Finally, we apply the method to identify SNP–SNP interactions for three phenotypes from the Alzheimer’s Disease Neuroimaging Initiative (ADNI), providing a real-world evaluation in a dataset comprising individuals of diverse ancestry.

## 2 Material and Methods

### 2.1 Analytical Calculation of Bias

We sought to quantify the spurious interaction effect due to population stratification — that is measure the bias of the estimated interaction term across regression models when samples are drawn from two distinct subpopulations and SNP effects differ between them, a setting commonly encountered in large-scale genomics studies (Mägi et al., 2017; Patel et al., 2022).

We considered a quantitative phenotype measured in unrelated individuals sampled from two populations, denoted A and B. The phenotype was assumed to depend on two biallelic SNPs, *S*_1_ and *S*_2_. For the *i*th individual, population membership was represented by *P_i_*, where *P_i_* = 0 indicated membership in population A and *P_i_* = 1 indicated membership in population B. Let *Y_i_* denote the quantitative phenotype, and let *S*_1_*_i_* and *S*_2_*_i_* denote the minor allele dosages, taking values in *{*0, 1, 2*}*, for the first and second SNP, respectively.

The population-specific mean of the phenotype was specified as

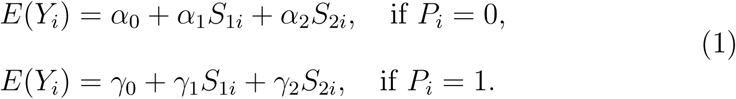

In this formulation, *α*_0_ denotes the intercept in population A, and *α*_1_ and *α*_2_ denote the corresponding main effects of SNPs *S*_1_ and *S*_2_. Similarly, *γ*_0_, *γ*_1_, and *γ*_2_ denote the intercept and SNP main effects in population B. Let *θ_jk_* denote the allele frequency of SNP *j* in population *k*. We further assumed that the linkage disequilibrium (LD) coefficient between *S*_1_ and *S*_2_ was *d*_1_ in population A and *d*_2_ in population B. Thus, both SNP effects, allele frequencies and the LD were allowed to vary across populations. No SNP-SNP interaction was assumed in either population.

The following three working regression models applied:

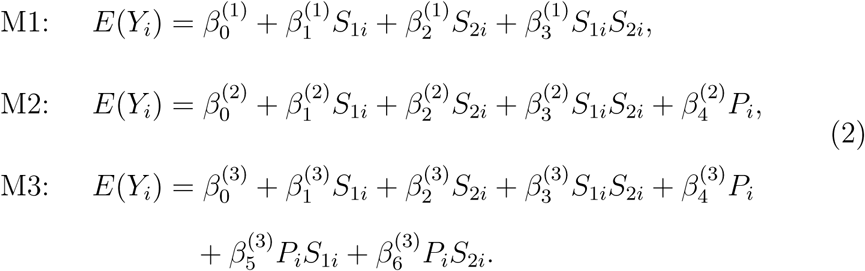

The first model (M1), we use a multiple regression framework that includes a multiplicative interaction term but does not adjust for population structure, as implemented in PLINK (Curtis et al., 2025; Purcell et al., 2007). In the second model (M2), we apply a commonly used strategy for correcting the inflation due to population structure adjustment in GWAS by incorporating population labels, in addition to the terms included in M1 an approach consistent with the EIGENSTRAT methodology. We found that the inflation in interaction test statistics could not be controlled by M2 which led us to develop a method (M3) based on the extension of the method developed by An et al. (2019) that incorporates the SNP-by-population substructure interaction term, alongside the terms used in M2. We then evaluated the impact of population structure by analytically deriving the bias of the estimated SNP-SNP interaction term *β*_3_ for the working regression model M2, denoted by *β^*_3_^(2)^.

The full analytical expressions, together with the derivations of the bias terms for models M2, and M3, are provided in the Supplementary Information. Since the true model does not include SNP-SNP interaction term, the bias of the interaction term for M2 can be written as

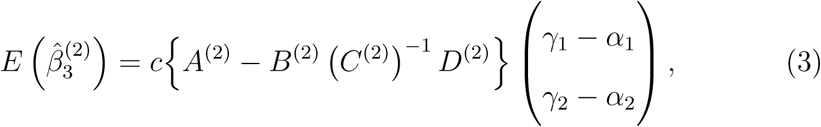

The complete expressions for *A*^(2)^*, B*^(2)^*, C*^(2)^*, D*^(2)^ and *c* can be found in the Appendix. The bias of the estimated SNP-SNP interaction coefficient was found to be proportional to the differences in population-specific parameters the SNP main effect differences (*γ*_1_ *− α*_1_) and (*γ*_2_ *− α*_2_). In contrast, the bias becomes zero under model M3, where both the population indicator *P_i_* and SNP-by-population interaction terms *S_j_ ×P* are included as covariates in the regression model because (1) is equivalent to M3 when *β*_3_^(2)^ as shown in the Supplementary Information.

It was noted that the bias of the interaction term becomes zero , if either of the two conditions are satisfied: (i) the main effects are same in both the populations for both the SNPs, (ii) the LD coefficients in both the populations are zero, along with no allele frequency differences among the SNPs in the two populations as demonstrated in the Supplementary Information.

The analytical expressions further show that increasing differences in minor allele frequencies between populations amplify the bias in the estimated interaction coefficient under M2. Differences in linkage disequilibrium between populations *d*_2_*−d*_1_, can also be a major source of bias in the interaction term.

### 2.2 Simulation studies

The impact of population structure on type I error rates was evaluated for both binary and continuous phenotypes using simulation studies. A total of 5,000 simulation replicates were performed for each phenotype type. Linear regression was used for continuous outcomes, whereas logistic regression was used for dichotomous outcomes.

Simulations were conducted under the four distinct scenarios with marginal effects but no interaction effects: (i) a single causal SNP in linkage equilibrium (LE, zero LD) with a second, non-causal SNP; (ii) a single causal SNP in linkage disequilibrium (random LD) with a second, non-causal SNP; (iii) two causal SNPs in LE; and (iv) two causal SNPs in LD.

For each simulation replicate, we considered two populations and generated genotype data for 1,000 individuals in total, with 500 individuals from each population and 400 SNPs per individual. Genotype and corresponding phenotype data were generated under varying levels of population differentiation, with Wright’s *F*_ST_ values set at 0.005, 0.01, and 0.02. These parameter choices are consistent with values reported in previous work (Cavalli-Sforza et al., 1993), which documented median *F*_ST_ estimates of approximately 0.008 among Europeans, 0.027 among Africans, and 0.01 among Asians.

Among these SNPs, the first two SNPs, *S*_1_ and *S*_2_, were considered to be bi-allelic loci with minor allele frequency greater than 0.05. These two SNPs were generated within the same haplotype block for scenarios (ii) and (iv), and independently for scenarios (i) and (iii), using the Balding–Nichols model (Balding & Nichols, 1995). For single main effect scenarios (i) and (ii), SNP *S*_2_ was assigned a zero main effect in scenarios.

For scenarios involving non-zero LD between *S*_1_ and *S*_2_, the ancestral haplotypes *h*_1_*, h*_2_*, h*_3_, and *h*_4_, corresponding to (*ab, aB, Ab, AB*), respectively, were assigned frequencies

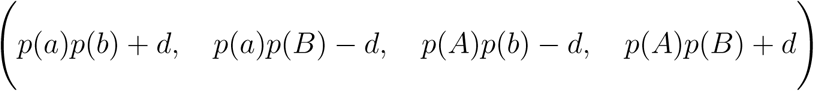

Here, *a* and *b* denote the minor alleles at *S*_1_ and *S*_2_, respectively, while *A* and *B* denote the corresponding major alleles. The ancestral minor allele frequencies *p*(*a*) and *p*(*b*) were independently sampled from a Uniform(0, 0.5) distribution. Conditional on these allele frequencies, the LD parameter *d* was sampled uniformly from its admissible bounds.

For each population, conditional on *h*_1_*, h*_2_*, h*_3_*, h*_4_ and *F*_ST_ haplotype frequencies were then sampled from a Dirichlet distribution with parameters

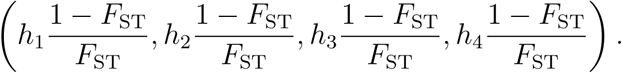

Given these population-specific haplotype frequencies, 2,000 haplotypes corresponding to 1,000 individuals were generated using multinomial sampling. For the remaining 398 SNPs, ancestral minor allele frequencies (*p*) were independently sampled from a Uniform(0, 0.5) distribution. Population-specific minor allele frequencies were subsequently drawn from a Beta distribution with parameters:

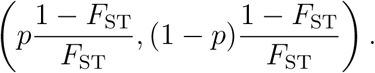

Genotypes for these SNPs were then assigned assuming Hardy–Weinberg equilibrium within each population.

For scenarios involving no LD between *S*_1_ and *S*_2_, all 400 SNPs were generated independently using the Balding–Nichols model. Across all four simulation scenarios, the minor allele frequency of the *j*th SNP in the *k*th population, denoted by *θ_jk_*, and the LD coefficients between the first two SNPs in the two populations, denoted by *d*_1_ and *d*_2_, were empirically calculated from the final genotype matrix *X*.

For the *i*th individual in the *k*th population, the continuous phenotype was generated by

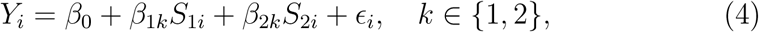

where *ɛ_i_ ∼ N* (0, 1).

For continuous phenotypes under scenarios (iii) and (iv), and for a fixed level of heritability, the main effect of SNP *S*_2_ in population A, denoted by *β*_21_, was determined while fixing the main effect of SNP *S*_1_ in population A at *β*_11_ = 0.10. Additionally, the difference in SNP effects between populations was held constant at *β*_diff_ = *β_j_*_1_ *− β_j_*_2_ = 0.35, for *j ∈ {*1, 2*}*.

Across all continuous phenotype scenarios, the proportion of individuals sampled from population A was fixed at *π* = 0.50. Heritability was varied across three levels, *h*^2^ *∈ {*0.1, 0.2, 0.3*}*. The exact expression used to compute *β*_21_ is provided in the Supplementary Information.

For the single main effect scenarios, only the difference in the main effect of SNP *S*_1_ across the two populations was fixed at *β*_diff_ = 0.35. The population-specific main effects of SNP *S*_1_ were then determined as functions of the minor allele frequencies in population A (*θ*_11_) and population B (*θ*_12_).

For dichotomous phenotypes, genotype data were generated for 4,000 individuals sampled from two populations. An underlying continuous latent phenotype was then simulated using the approach described above. Individuals whose latent phenotype exceeded a prespecified threshold were classified as cases, whereas the remaining individuals were classified as controls. The threshold was chosen to yield a population disease prevalence of 0.20. After the assignment, 500 cases and 500 controls were randomly selected to reflect the sampling in case-control studies.

### 2.3 Data Analysis Models

For continuous phenotypes, models M1, M2, and M3 were fitted using linear regression. Because true population membership, *P_i_*, is generally unobserved in practice, population structure was represented using the top 10 principal components *PC_li_* (*l* = 1, 2*, . . . ,* 10) derived from the genetic relatedness matrix *W* . The genetic relatedness matrix was defined as

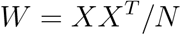

where *X*denotes the standardized genotype matrix, with each column corresponding to a SNP and *N* denotes the total number of SNPs in our dataset. The corresponding working models are as follows:

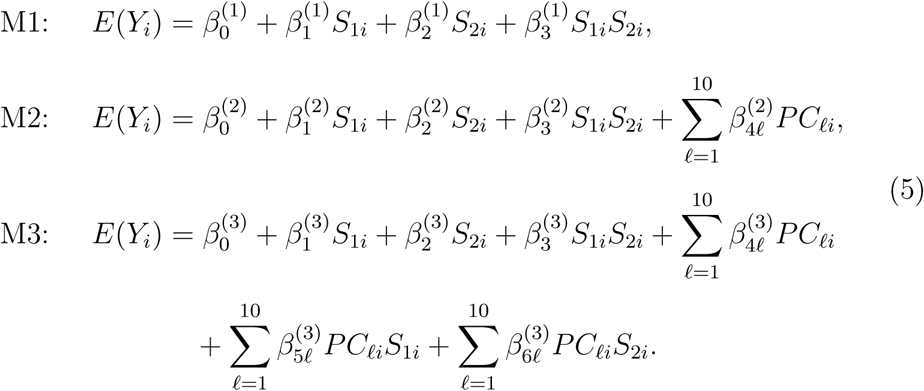

For dichotomous phenotypes, the corresponding models were fitted using logistic regression, and the significance of the interaction term was assessed using the Wald test or likelihood ratio test (LRT).

When fitting M3 for both continuous and dichotomous phenotypes, we frequently observed multicollinearity among predictors, which led to inflated false positive rates. To mitigate this issue, SNP-by-principal-component interaction terms (*PC_ℓ_S*_1_*, PC_ℓ_S*_2_) in M3 with a variance inflation factor (VIF) exceeding 100 were removed from the model.

### 2.4 Real data analysis

We applied our proposed method to the ADNI-GWAS dataset obtained from the publicly available data of the Alzheimer’s Disease Neuroimage Initiative (ADNI) database (adni.loni.usc.edu). The ADNI was launched in 2003 as a public-private partnership, led by Principal Investigator Michael W. Weiner, MD. The primary goal of ADNI has been to test whether serial magnetic resonance imaging (MRI), positron emission tomography (PET), other biological markers, and clinical and neuropsychological assessments can be combined to measure the progression of mild cognitive impairment (MCI) and early Alzheimer’s disease (AD). For up-to-date information, see www.adniinfo.org. ADNI is an ongoing, longitudinal study with primary purpose being to explore the genetic and neuroimaging information associated with lateonset Alzheimer’s disease (LOAD). The study investigators recruited elderly subjects older than 65 years of age comprising about 400 subjects with mild cognitive impairment (MCI), about 200 subjects with Alzheimer’s disease (AD), and about 200 healthy controls. Each subject was followed for at least 3 years. During the study period, the subjects were assessed with magnetic resonance imaging (MRI) measures and psychiatric evaluation to determine the diagnosis status at each time point.

GWAS genotype data were generated from genomic DNA extracted from peripheral blood samples of ADNI-3 participants using two Illumina genotyping arrays: the Illumina Infinium Global Screening Array v2 (GSA2) for 327 participants and the Illumina Infinium Global Screening Array-24 v3 Bead-Chip for 328 participants. The two datasets were merged in a single file. See Saykin et al. (2015) for more information. The data initially included 792,113 SNPs. We excluded one individual from each pair with evidence of cryptic relatedness, defined as a PLINK pairwise *π*^ statistic greater than 0.25 (Purcell et al., 2007). We then applied further quality control measures by excluding SNPs with missing genotype rate *>* 0.1, Hardy–Weinberg equilibrium test *p*-value *<* 10*^−^*^7^, and minor allele frequency *<* 5%; the total number of remaining SNPs was 481,475 for 648 individuals and 6,955 on chromosome 22.

All pairwise epistasis analyses on the 6,955 SNPs were performed for three phenotypes: whole-brain volume (Whole Brain), hippocampal volume (Hippocampus), and Mini-Mental State Examination (MMSE) score. Wholebrain volume and hippocampal volume were treated as continuous phenotypes, whereas MMSE score, although ordinal in nature, was analyzed as a continuous phenotype in the primary analysis. After removing individuals with missing phenotypes, we were left with 635 individuals for MMSE, 527 for Hippocampus and 548 for Whole Brain. For each phenotype, we compared the three regression models using linear regression with adjustment for the top 10 principal components generated from the autosomal SNPs.

To evaluate the performance of logistic regression, MMSE scores were additionally dichotomized using a threshold of 26 (Kvitting et al., 2019).

## 3 Results

### 3.1 Simulation Results

Table 1 shows the type I error rates for the three different models (M1, M2, M3) for quantitative phenotypes where the significance of the interaction term is tested using *F* -test. Supplementary Figure S1-S4 shows the Q-Q plots corresponding to the *p*-values obtained from the tests. We observed an inflation of interaction test statistics across all four simulation scenarios described in the methods section. Overall, the inflation was highest in the case where both SNPs were associated with the phenotype, along with highest heritability and highest population differentiation among the two populations(high *F*_ST_). While in this case M2 performed slightly better than M1, it could not control the false positive rates — which was significantly better controlled by M3. Inflation was smallest in scenario (i), in which only one SNP was associated with the phenotype, while the second SNP was independent of both the phenotype and the causal SNP. In this setting, M3 generally performed best among the three models; however, M1 outperformed M2 in some instances.

**Table 1:** Type I error for continuous phenotypes at significance level of 0.05.

| Case | $F_{ST}$ | $h^2$ | M1 | M2 | M3 |
| --- | --- | --- | --- | --- | --- |
| 1snp zero ld | 0.005 | 0.1 | 0.0634 | 0.0598 | 0.0556 |
|  | 0.005 | 0.2 | 0.0596 | 0.0590 | 0.0546 |
|  | 0.005 | 0.3 | 0.0584 | 0.0598 | 0.0512 |
|  | 0.010 | 0.1 | 0.0680 | 0.0692 | 0.0540 |
|  | 0.010 | 0.2 | 0.0754 | 0.0762 | 0.0528 |
|  | 0.010 | 0.3 | 0.0690 | 0.0700 | 0.0512 |
|  | 0.020 | 0.1 | 0.0734 | 0.0758 | 0.0472 |
|  | 0.020 | 0.2 | 0.0724 | 0.0734 | 0.0470 |
|  | 0.020 | 0.3 | 0.0764 | 0.0784 | 0.0498 |
| 1snp random ld | 0.005 | 0.1 | 0.0688 | 0.0648 | 0.0520 |
|  | 0.005 | 0.2 | 0.0642 | 0.0596 | 0.0508 |
|  | 0.005 | 0.3 | 0.0670 | 0.0642 | 0.0574 |
|  | 0.010 | 0.1 | 0.0766 | 0.0688 | 0.0550 |
|  | 0.010 | 0.2 | 0.0766 | 0.0714 | 0.0538 |
|  | 0.010 | 0.3 | 0.0756 | 0.0682 | 0.0570 |
|  | 0.020 | 0.1 | 0.0898 | 0.0764 | 0.0482 |
|  | 0.020 | 0.2 | 0.0912 | 0.0754 | 0.0518 |
|  | 0.020 | 0.3 | 0.0892 | 0.0736 | 0.0458 |
| 2snp zero ld | 0.005 | 0.1 | 0.0680 | 0.0648 | 0.0456 |
|  | 0.005 | 0.2 | 0.0646 | 0.0664 | 0.0508 |
|  | 0.005 | 0.3 | 0.0648 | 0.0620 | 0.0506 |
|  | 0.010 | 0.1 | 0.0812 | 0.0746 | 0.0498 |
|  | 0.010 | 0.2 | 0.0842 | 0.0796 | 0.0508 |
|  | 0.010 | 0.3 | 0.0742 | 0.0686 | 0.0508 |
|  | 0.020 | 0.1 | 0.0928 | 0.0898 | 0.0514 |
|  | 0.020 | 0.2 | 0.0986 | 0.0956 | 0.0542 |
|  | 0.020 | 0.3 | 0.0876 | 0.0866 | 0.0504 |
| 2snp random ld | 0.005 | 0.1 | 0.0998 | 0.0848 | 0.0620 |
|  | 0.005 | 0.2 | 0.1030 | 0.0882 | 0.0608 |
|  | 0.005 | 0.3 | 0.1002 | 0.0836 | 0.0616 |
|  | 0.010 | 0.1 | 0.1300 | 0.0962 | 0.0554 |
|  | 0.010 | 0.2 | 0.1270 | 0.0938 | 0.0540 |
|  | 0.010 | 0.3 | 0.1322 | 0.0992 | 0.0540 |
|  | 0.020 | 0.1 | 0.1676 | 0.1222 | 0.0568 |
|  | 0.020 | 0.2 | 0.1690 | 0.1194 | 0.0458 |
|  | 0.020 | 0.3 | 0.1716 | 0.1194 | 0.0512 |

*Note:* Empirical type I error rates are reported for the SNP-SNP interaction term under three regression models, M1, M2, and M3, using an $F$ -test at a significance level of 0.05. For each combination of $F_{ST}$ and heritability, 5,000 simulation replicates were performed. The scenario “1snp zero ld” denotes the setting in which only one SNP has a non-zero main effect and is independent of a second SNP with zero main effect. The scenario “1snp random ld” denotes the setting in which one SNP with a non-zero main effect is in random LD with a second SNP with zero main effect. The scenario “2snp zero ld” denotes the setting in which two independent SNPs both have non-zero main effects. The scenario “2snp random ld” denotes the setting in which both SNPs have non-zero main effects and are in random LD.

Similar to the case of continuous phenotypes, the inflation of interaction test statistics for binary phenotypes was highest in case (iv), and the least in case (i). M3 performed better at false positive correction than M2 and M1 in all the cases for Wald test, but the same could not be said for LRT. Overall the M3 of Wald test performed better than LRT in most of the cases. The type I error rates for Wald test are shown in Table 2 and the corresponding Q-Q plots are shown in Supplementary Figure S5-S8. The type I error rates for LRT are shown in Supplementary Table S1 and the corresponding Q-Q plots are shown in Supplementary Figure S9-S12. While performing M3 logistic regression, we often encountered infinite estimates of *β*_3_ due to certain SNP-by-principal-component interaction terms being collinear with other covariates. Thus we removed all such columns which had a VIF of more than 100.

**Table 2:** Type I error for dichotomous phenotypes at significance level of 0.05.

| Case | $F_{ST}$ | $h^2$ | M1 | M2 | M3 |
| --- | --- | --- | --- | --- | --- |
| 1snp zero ld | 0.005 | 0.1 | 0.0464 | 0.0474 | 0.0544 |
|  | 0.005 | 0.2 | 0.0464 | 0.0472 | 0.0534 |
|  | 0.005 | 0.3 | 0.0378 | 0.0368 | 0.0458 |
|  | 0.010 | 0.1 | 0.0530 | 0.0538 | 0.0512 |
|  | 0.010 | 0.2 | 0.0508 | 0.0540 | 0.0490 |
|  | 0.010 | 0.3 | 0.0536 | 0.0552 | 0.0590 |
|  | 0.020 | 0.1 | 0.0594 | 0.0614 | 0.0596 |
|  | 0.020 | 0.2 | 0.0558 | 0.0550 | 0.0508 |
|  | 0.020 | 0.3 | 0.0484 | 0.0506 | 0.0434 |
| 1snp random ld | 0.005 | 0.1 | 0.0542 | 0.0536 | 0.0614 |
|  | 0.005 | 0.2 | 0.0594 | 0.0570 | 0.0616 |
|  | 0.005 | 0.3 | 0.0466 | 0.0476 | 0.0492 |
|  | 0.010 | 0.1 | 0.0588 | 0.0566 | 0.0574 |
|  | 0.010 | 0.2 | 0.0570 | 0.0554 | 0.0544 |
|  | 0.010 | 0.3 | 0.0500 | 0.0492 | 0.0478 |
|  | 0.020 | 0.1 | 0.0750 | 0.0698 | 0.0610 |
|  | 0.020 | 0.2 | 0.0748 | 0.0656 | 0.0590 |
|  | 0.020 | 0.3 | 0.0622 | 0.0554 | 0.0526 |
| 2snp zero ld | 0.005 | 0.1 | 0.0592 | 0.0584 | 0.0560 |
|  | 0.005 | 0.2 | 0.0898 | 0.0892 | 0.0600 |
|  | 0.005 | 0.3 | 0.1052 | 0.1062 | 0.0636 |
|  | 0.010 | 0.1 | 0.0740 | 0.0706 | 0.0596 |
|  | 0.010 | 0.2 | 0.1010 | 0.1002 | 0.0684 |
|  | 0.010 | 0.3 | 0.1098 | 0.1150 | 0.0696 |
|  | 0.020 | 0.1 | 0.0842 | 0.0846 | 0.0556 |
|  | 0.020 | 0.2 | 0.1044 | 0.1046 | 0.0582 |
|  | 0.020 | 0.3 | 0.1142 | 0.1174 | 0.0640 |
| 2snp random ld | 0.005 | 0.1 | 0.0850 | 0.0764 | 0.0622 |
|  | 0.005 | 0.2 | 0.1024 | 0.0918 | 0.0744 |
|  | 0.005 | 0.3 | 0.1098 | 0.1068 | 0.0644 |
|  | 0.010 | 0.1 | 0.1036 | 0.0826 | 0.0616 |
|  | 0.010 | 0.2 | 0.1264 | 0.1002 | 0.0718 |
|  | 0.010 | 0.3 | 0.1138 | 0.1104 | 0.0666 |
|  | 0.020 | 0.1 | 0.1366 | 0.1084 | 0.0556 |
|  | 0.020 | 0.2 | 0.1450 | 0.1174 | 0.0568 |
|  | 0.020 | 0.3 | 0.1312 | 0.1150 | 0.0674 |

*Note:* Empirical type I error rates are reported for the SNP-SNP interaction term under three regression models, M1, M2, and M3, using the Wald test at a significance level of 0.05. For each combination of $F_{ST}$ and heritability, 5,000 simulation replicates were performed. The scenario “1snp zero ld” denotes the setting in which only one SNP has a non-zero main effect and is independent of a second SNP with zero main effect. The scenario “1snp random ld” denotes the setting in which one SNP with a non-zero main effect is in random LD with a second SNP with zero main effect. The scenario “2snp zero ld” denotes the setting in which two independent SNPs both have non-zero main effects. The scenario “2snp random ld” denotes the setting in which both SNPs have non-zero main effects and are in random LD.

### 3.2 Real Data Analysis Results

We performed epistasis analyses of common variants for three continuous phenotypes obtained from ADNI: MMSE score, hippocampal volume, and whole-brain volume. For each phenotype, we used linear regression and *F* tests to compare the proportion of significant SNP-SNP interactions across different significance levels under models M1, M2, and M3. The results are presented in Table 3, with the corresponding Q-Q plots shown in Figure 1. Among the three phenotypes, MMSE showed the clearest reduction in inflation of interaction test *p*-values under M3, whille M1 and M2 showed inflation especially at *α* = 5 *×* 10*^−^*^6^. Interaction test statistics for both Whole Brain and Hippocampus showed minimal inflation, however the proportion of significant SNP-SNP interactions for Whole Brain at *α* = 5 *×* 10*^−^*^6^ was slightly higher for M1 compared to M2 and M3.

**Figure 1:**
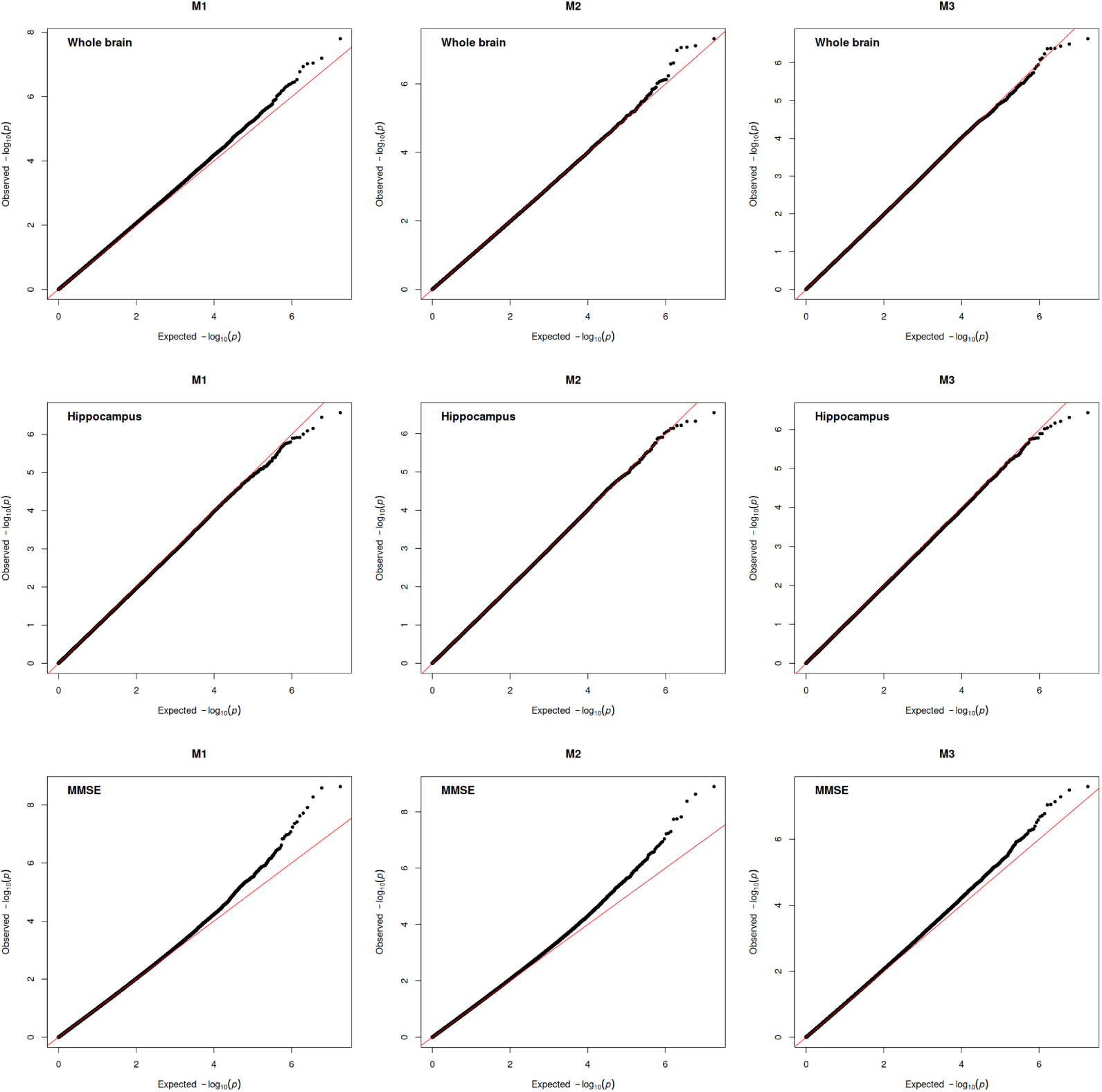
Q-Q plots of *p*-values from *F* -test of the SNP-SNP interaction term under linear regression models M1, M2, and M3, for three continuous phenotypes: Whole Brain , Hippocampus and MMSE scores

**Table 3:** ADNI data analysis.

| Phenotype | $\alpha$ | M1 | M2 | M3 |
| --- | --- | --- | --- | --- |
| WholeBrain | $5 \times 10^{-2}$ | $5.41 \times 10^{-2}$ | $4.83 \times 10^{-2}$ | $4.86 \times 10^{-2}$ |
| | $5 \times 10^{-4}$ | $6.49 \times 10^{-4}$ | $4.93 \times 10^{-4}$ | $5.04 \times 10^{-4}$ |
| | $5 \times 10^{-6}$ | $8.72 \times 10^{-6}$ | $5.44 \times 10^{-6}$ | $3.97 \times 10^{-6}$ |
| Hippocampus | $5 \times 10^{-2}$ | $4.71 \times 10^{-2}$ | $4.84 \times 10^{-2}$ | $4.76 \times 10^{-2}$ |
| | $5 \times 10^{-4}$ | $4.53 \times 10^{-4}$ | $4.89 \times 10^{-4}$ | $4.49 \times 10^{-4}$ |
| | $5 \times 10^{-6}$ | $3.30 \times 10^{-6}$ | $4.66 \times 10^{-6}$ | $4.32 \times 10^{-6}$ |
| MMSE | $5 \times 10^{-2}$ | $5.00 \times 10^{-2}$ | $5.28 \times 10^{-2}$ | $5.30 \times 10^{-2}$ |
| | $5 \times 10^{-4}$ | $6.32 \times 10^{-4}$ | $7.07 \times 10^{-4}$ | $6.94 \times 10^{-4}$ |
| | $5 \times 10^{-6}$ | $1.66 \times 10^{-5}$ | $1.88 \times 10^{-5}$ | $9.99 \times 10^{-6}$ |

*Note:* Proportion of significant SNP-SNP interactions for three phenotypes: Whole-Brain, Hippocampus, and MMSE scores. Results are shown under three regression models (M1, M2, and M3) and three significance thresholds: $5 \times 10^{-2}$ , $5 \times 10^{-4}$ , and $5 \times 10^{-6}$ .

We further dichotomized MMSE using a cutoff of 26, assigning individuals with scores greater than 26 as controls and those with scores of 26 or lower as cases. The corresponding proportions and Q-Q plots are presented in Table 4 and Figure 2, respectively. We observed that the *p*-values at different significance levels for M3 was closer to nominal significance levels compared to M1 and M2, where M1 and M2 gave deflated *p*-values. SNP-SNP interactions which resulted in infinite estimate of regression coefficient were excluded from subsequent analysis.

**Figure 2:**
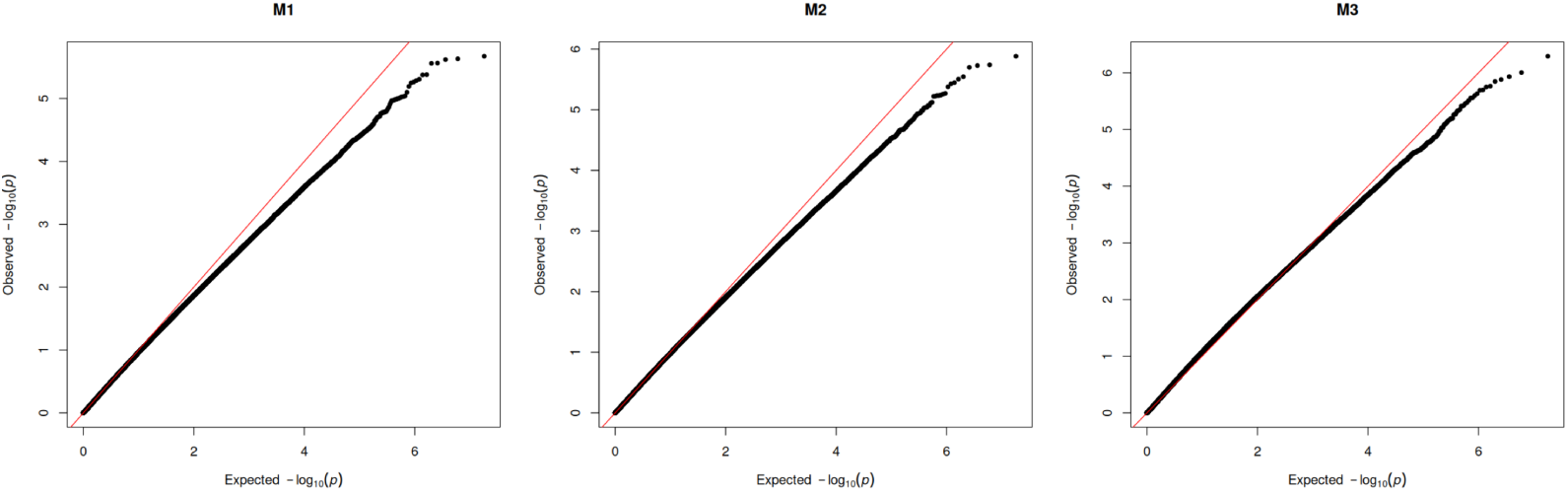
Q-Q plots of *p*-values from Wald tests of the SNP-SNP interaction term under logistic regression models M1, M2, and M3, with dichotomized MMSE (cutoff of 26) score as the phenotype.

**Table 4:** ADNI data Analysis (Binary)

| Phenotype | $\alpha$ | M1 | M2 | M3 |
| --- | --- | --- | --- | --- |
| MMSE (binary) | $5 \times 10^{-2}$ | $4.26 \times 10^{-2}$ | $4.58 \times 10^{-2}$ | $5.99 \times 10^{-2}$ |
| | $5 \times 10^{-4}$ | $2.20 \times 10^{-4}$ | $2.67 \times 10^{-4}$ | $4.12 \times 10^{-4}$ |
| | $5 \times 10^{-6}$ | $8.88 \times 10^{-7}$ | $9.99 \times 10^{-7}$ | $2.55 \times 10^{-6}$ |

*Note.* Proportion of significant SNP-SNP interactions for dichotomized MMSE phenotype based on the cutoff of 26. Results are shown under three regression models (M1, M2, and M3) and three significance thresholds: $5 \times 10^{-2}$ , $5 \times 10^{-4}$ , and $5 \times 10^{-6}$ .

## 4 Discussion

In this study, we examined the impact of population stratification on epistasis detection when samples are ascertained from two distinct ancestral populations. We propose a method whereby including SNP-by-principal-component interaction terms, reduces false positive inflation of SNP-SNP interaction test statistics arising from population-specific differences in allele frequencies, linkage disequilibrium patterns, and SNP main effects. Our results suggest that the reproducibility of gene-gene interaction findings may be improved by more appropriately accounting for population structure, as implemented in the proposed model.

Several limitations should be noted. First, our approach uses principal components as proxies for true population labels. Although principal components are widely used to adjust for global ancestry, they may not fully capture local ancestry or population structure that varies across genomic regions, particularly in admixed populations. As a result, residual confounding may persist when local ancestry effects are present (An et al., 2019).

Second, the proposed approach was less effective in controlling false positives when rare variants were involved, particularly when at least one of the SNPs with a population-specific main effect was rare. In our analyses, Firth regression (Firth, 1993) appeared to provide better control in such settings than the Wald test or likelihood ratio test. However, the computational burden of Firth regression makes its routine application to genome-wide epistasis scans challenging. Future work should therefore focus on improving the computational efficiency of Firth-based estimation and testing procedures for rare-variant interaction analyses.

Third, multicollinearity among SNP-by-principal-component interaction terms posed a practical challenge in model fitting. In this study, we addressed this issue by removing interaction terms with a VIF greater than 100. Although this strategy reduced estimation instability, further work is needed to determine the cutoff for selecting which covariates should be retained or excluded from the model.

Finally, our analysis focused on additive two-SNP models in which SNP effects were represented by allele dosage. We did not consider alternative genetic models, such as dominant or recessive coding of SNP effects. Extending the proposed framework to accommodate non-additive genotype coding schemes as examined by Ueki and Cordell (2012), as well as more complex ancestry structures, represents an important direction for future research.

## Data availability statement

The authors do not own data used in the manuscript. Data obtained were collected and owned by the Alzheimer’s Disease Neuroimaging Initiative (ADNI). Researchers may request and access the data through the ADNI website (http://adni.loni.usc.edu/). The authors had no special access privileges to this data.

## Declaration of AI usage

During the preparation of this work the authors used publicly available AI models ChatGPT(5.5) and Gemini(3.1,3.5) in order to improve readability and language. After using this tool, the authors reviewed and edited the content as needed and take full responsibility for the content of the published article.

## Acknowledgements

Data collection and sharing for this project was funded by the Alzheimer’s Disease Neuroimaging Initiative (ADNI) (National Institutes of Health Grant U01 AG024904) and DOD ADNI (Department of Defense award number W81XWH-12-2-0012). ADNI is funded by the National Institute on Aging, the National Institute of Biomedical Imaging and Bioengineering, and through generous contributions from the following: AbbVie, Alzheimer’s Association; Alzheimer’s Drug Discovery Foundation; Araclon Biotech; BioClinica, Inc.; Biogen; Bristol-Myers Squibb Company; CereSpir, Inc.; Eisai Inc.; Elan Pharmaceuticals, Inc.; EliLilly and Company; EuroImmun; F. Hoffmann-La Roche Ltd and its affiliated company Genentech, Inc.; Fujirebio; GE Healthcare; IXICO Ltd.; Janssen Alzheimer Immunotherapy Research & Development, LLC.; Johnson & Johnson Pharmaceutical Research & Development LLC.; Lumosity; Lundbeck; Merck & Co., Inc.; Meso Scale Diagnostics, LLC.; NeuroRx Research; Neurotrack Technologies; Novartis Pharmaceuticals Corporation; Pfizer Inc.; Piramal Imaging; Servier; Takeda Pharmaceutical Company; and Transition Therapeutics. The Canadian Institutes of Health Research is providing funds to support ADNI clinical sites in Canada. Private sector contributions are facilitated by the Foundation for the National Institutes of Health (www.fnih.org). The grantee organization is the Northern California Institute for Research and Education, and the study is coordinated by the Alzheimer’s Disease Cooperative Study at the University of California, San Diego. ADNI data are disseminated by the Laboratory for Neuro Imaging at the University of Southern California.

This work was partially supported by JSPS KAKENHI Grant Number 26K14742 and JST SPRING, Japan Grant Number JPMJSP2172.

## Appendix

We write the expressions of the variables in Equation (3).

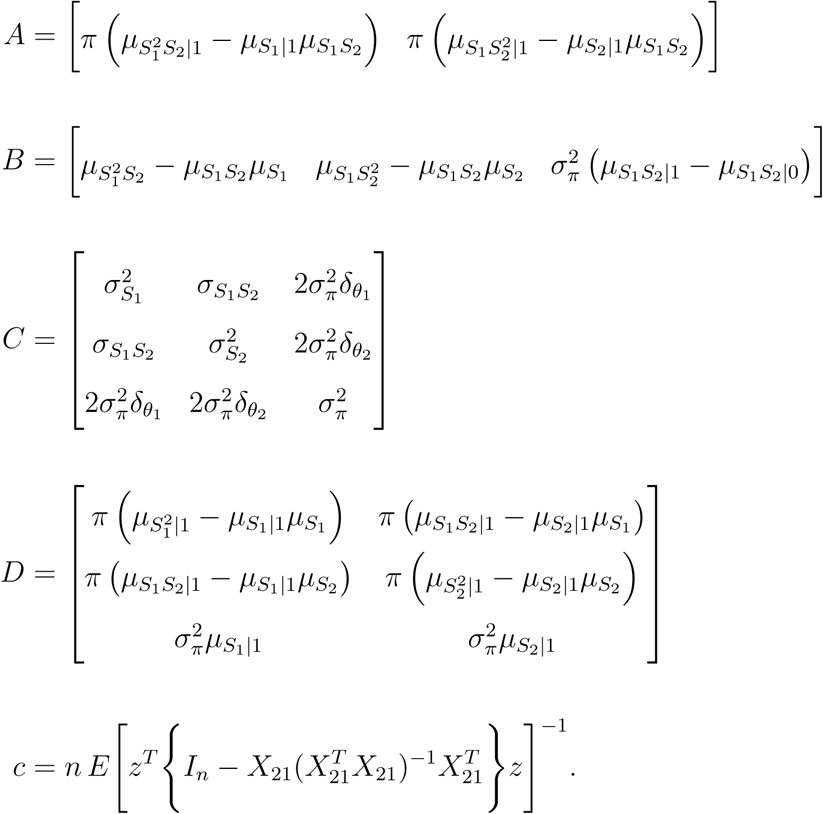

Here, *z* denotes the *n ×* 1 vector of centered SNP-SNP interaction terms, and *I_n_* denotes the *n × n* identity matrix. The matrix *X*_21_ is an *n ×* 3 design matrix whose first, second, and third columns contain the centered values of *S*_1_, *S*_2_, and the population label *P* , respectively. The notation *µ_f_*_(_*_S_*_1_*_,S_*_2)_*_|k_* represents the conditional expectation of a function *f* of *S*_1_ and *S*_2_, *f* (*S*_1_*, S*_2_), within *k*th population , whereas *µ_f_*_(_*_S_*_1_*_,S_*_2)_ denotes the corresponding marginal expectation. Similarly, *σ*^2^ denotes the marginal variance of *S_j_*, and *σ_S_*_1_*_S_*_2_ denotes the covariance between *S*_1_ and *S*_2_. The term *σ*^2^ represents the sampling variance of individuals across populations, and *δ_θj_* denotes the populationlevel difference in the minor allele frequency of the *j*th SNP. Throughout, the constant *c* is positive. Detailed expressions are given in the Supplementary Information.

## References

1. Abdellaoui, A., Yengo, L., Verweij, K. J. H., & Visscher, P. M. (2023). 15 years of GWAS discovery: Realizing the promise. American Journal of Human Genetics, 110 (2), 179–194. 10.1016/j.ajhg.2022.12.011

2. Abegaz, F., Van Lishout, F., Mahachie John, J. M., Chiachoompu, K., Bhardwaj, A., Duroux, D., Gusareva, E. S., Wei, Z., Hakonarson, H., & Van Steen, K. (2021). Performance of model-based multifactor dimensionality reduction methods for epistasis detection by controlling population structure. BioData Mining, 14, 16. 10.1186/s13040-021-00247-w

3. An, J., Won, S., Hecker, J., & Lange, C. (2019). Effect of population stratification on SNP-by-environment interaction. Genetic Epidemiology, 43, 1046–1055. 10.1002/gepi.22250

4. Balding, D. J., & Nichols, R. A. (1995). A method for quantifying differentiation between populations at multi-allelic loci and its implications for investigating identity and paternity. Genetica, 96 (1–2), 3–12. 10.1007/BF01441146

5. Cavalli-Sforza, L. L., Menozzi, P., & Piazza, A. (1993). Demic expansions and human evolution. Science, 259 (5095), 639–646. 10.1126/science.8430313

6. Cordell, H. J. (2002). Epistasis: What it means, what it doesn’t mean, and statistical methods to detect it in humans. Human Molecular Genetics, 11 (20), 2463–2468. 10.1093/hmg/11.20.2463

7. Cordell, H. J. (2009). Detecting gene-gene interactions that underlie human diseases. Nature Reviews Genetics, 10 (6), 392–404. 10.1038/nrg2579

8. Curtis, A. A., Yu, Y., Carey, M., Yilmaz, Y. E., & Savas, S. (2025). A genomewide SNP-SNP interaction analysis exploring novel interacting loci associated with the risk of recurrence in colorectal cancer. PLOS ONE, 20 (6), e0321967. 10.1371/journal.pone.0321967

9. Devlin, B., & Roeder, K. (1999). Genomic control for association studies. Biometrics, 55 (4), 997–1004. 10.1111/j.0006-341X.1999.00997.x

10. Firth, D. (1993). Bias reduction of maximum likelihood estimates. Biometrika, 80 (1), 27–38. 10.2307/2336755

11. Goudey, B., Abraham, G., Kikianty, E., Wang, Q., Rawlinson, D., Shi, F., Haviv, I., Stern, L., Kowalczyk, A., & Inouye, M. (2017). Interactions within the MHC contribute to the genetic architecture of celiac disease. PLOS ONE, 12 (3), e0172826. 10.1371/journal.pone.0172826

12. Hellwege, J. N., Keaton, J. M., Giri, A., Gao, X., Velez Edwards, D. R., & Edwards, T. L. (2017). Population stratification in genetic association studies. Current Protocols in Human Genetics, 95 (1), 1.22.1–1.22.23. 10.1002/cphg.48

13. Jiang, L., Zheng, Z., Qi, T., Kemper, K. E., Wray, N. R., Visscher, P. M., & Yang, J. (2019). A resource-efficient tool for mixed model association analysis of large-scale data. Nature Genetics, 51 (12), 1749–1755. 10.1038/s41588-019-0530-8

14. Kang, H. M., Sul, J. H., Service, S. K., Zaitlen, N. A., Kong, S.-y., Freimer, N. B., Sabatti, C., & Eskin, E. (2010). Variance component model to account for sample structure in genome-wide association studies. Nature Genetics, 42 (4), 348–354. 10.1038/ng.548

15. Kvitting, A. S., Fällman, K., Wressle, E., & Marcusson, J. (2019). Age-normative MMSE data for older persons aged 85 to 93 in a longitudinal swedish cohort. Journal of the American Geriatrics Society, 67 (3), 534–538. 10.1111/jgs.15694

16. Mägi, R., Horikoshi, M., Sofer, T., Mahajan, A., Kitajima, H., Franceschini, N., McCarthy, M. I., COGENT-Kidney Consortium, T2D-GENES Consortium, & Morris, A. P. (2017). Trans-ethnic meta-regression of genome-wide association studies accounting for ancestry increases power for discovery and improves fine-mapping resolution. Human Molecular Genetics, 26 (18), 3639–3650. 10.1093/hmg/ddx280

17. Ning, C., Wang, D., Kang, H., Mrode, R., Zhou, L., Xu, S., & Liu, J.-F. (2018). A rapid epistatic mixed-model association analysis by linear retransformations of genomic estimated values. Bioinformatics, 34 (11), 1817–1825. 10.1093/bioinformatics/bty017

18. Niu, A., Zhang, S., & Sha, Q. (2011). A novel method to detect gene–gene interactions in structured populations: MDR-SP. Annals of Human Genetics, 75 (6), 742–754. 10.1111/j.1469-1809.2011.00681.x

19. Patel, R. A., Musharoff, S. A., Spence, J. P., Pimentel, H., Tcheandjieu, C., Mostafavi, H., Sinnott-Armstrong, N., Clarke, S. L., Smith, C. J., V.A. Million Veteran Program, Durda, P. P., Taylor, K. D., Tracy, R., Liu, Y., Johnson, W. C., Aguet, F., Ardlie, K. G., Gabriel, S., Smith, J., . . . Pritchard, J. K. (2022). Genetic interactions drive heterogeneity in causal variant effect sizes for gene expression and complex traits. American Journal of Human Genetics, 109 (7), 1286–1297. 10.1016/j.ajhg.2022.05.014

20. Price, A. L., Patterson, N. J., Plenge, R. M., Weinblatt, M. E., Shadick, N. A., & Reich, D. (2006). Principal components analysis corrects for stratification in genome-wide association studies. Nature Genetics, 38 (8), 904–909. 10.1038/ng1847

21. Purcell, S., Neale, B., Todd-Brown, K., Thomas, L., Ferreira, M. A. R., Bender, D., Maller, J., Sklar, P., de Bakker, P. I. W., Daly, M. J., & Sham, P. C. (2007). PLINK: A tool set for whole-genome association and population-based linkage analyses. American Journal of Human Genetics, 81 (3), 559–575. 10.1086/519795

22. Saykin, A. J., Shen, L., Yao, X., Kim, S., Nho, K., Risacher, S. L., Ramanan, V. K., Foroud, T. M., Faber, K. M., Sarwar, N., Munsie, L. M., Hu, X., Soares, H. D., Potkin, S. G., Thompson, P. M., Kauwe, J. S. K., Kaddurah-Daouk, R., Green, R. C., Toga, A. W., & Weiner, M. W. (2015). Genetic studies of quantitative MCI and AD phenotypes in ADNI: Progress, opportunities, and plans. Alzheimer’s & Dementia, 11 (7), 792–814. 10.1016/j.jalz.2015.05.009

23. Tyler, A. L., El Kassaby, B., Kolishovski, G., Emerson, J., Wells, A. E., Mahoney, J. M., & Carter, G. W. (2021). Effects of kinship correction on inflation of genetic interaction statistics in commonly used mouse populations. G3: Genes, Genomes, Genetics, 11 (7), jkab131. 10.1093/g3journal/jkab131

24. Ueki, M., & Cordell, H. J. (2012). Improved statistics for genome-wide interaction analysis. PLOS Genetics, 8 (4), e1002625. 10.1371/journal.pgen.1002625

